# The conundrum of Shiga toxin-producing *Escherichia coli* O157:H7 persistence: Evidence for locally persistent lineages

**DOI:** 10.1101/2025.09.29.679259

**Authors:** Gillian A.M. Tarr, William Finical, Joshua M. Rounds, Anna Panek, Kirk Smith

**Affiliations:** Division of Environmental Health Sciences, School of Public Health, University of Minnesota, Minneapolis, MN, 55455, U.S.A; Foodborne, Waterborne, Vectorborne, and Zoonotic Diseases Section, Minnesota Department of Health, St. Paul, MN, 55164, U.S.A; Division of Epidemiology and Community Health, School of Public Health, University of Minnesota, Minneapolis, MN, 55455, U.S.A

**Keywords:** Shiga toxin-producing *Escherichia coli*, disease ecology, phylogenetics, persistence, genomic epidemiology

## Abstract

Evidence suggests that Shiga toxin-producing *Escherichia coli* (STEC) strains do not persist at the farm level. We hypothesized that ecosystem-level STEC persistence occurs and contributes significantly to disease burden. We tested this by identifying locally persistent lineages (LPLs) of STEC O157:H7 in Minnesota. We identified 15 distinct LPLs, which were associated with 35.3% of reported cases in Minnesota and persisted for 1.3 to 8.6 years. LPLs were associated with multiple outbreaks with Minnesota sources and no multi-state outbreaks, and LPL cases were spatially clustered. Our findings show long-term persistence in defined geographic areas, suggesting the importance of ecosystem-level persistence.

**Summary:** Shiga toxin-producing *Escherichia coli* (STEC) persistence is poorly understood. This study examined long-term STEC persistence. Locally persistent lineages (LPLs) of STEC O157:H7 contributed significantly to disease burden and persisted up to 8.6 years. LPLs suggest the importance of ecosystem-level persistence.

## Background

Nationally persistent strains of Shiga toxin-producing *Escherichia coli* (STEC) O157:H7 have been associated with several high-profile outbreaks.^1^ However, the contribution of nationally persistent strains to STEC disease burden may be dwarfed by locally persistent strains. In Alberta, Canada, which has an exceptionally large cattle industry and high STEC burden, strains persisting in the province for up to 13 years were associated with 75% of recently reported STEC O157 cases.^2^ We termed these strains locally persistent lineages (LPLs) to distinguish them from strains persisting through regional, national, or international migration, as local persistence implies a local reservoir.

However, the outsized role of LPLs in STEC O157 disease burden in Alberta may be more the exception than the rule. Ruminants, particularly cattle, are considered STEC’s primary reservoir,^3^ and several lines of evidence suggest that STEC are not maintained long-term in ruminant populations. Most cattle are colonized <60 days, and reinfections are often cleared within a week.^4^ At the farm level, large fluctuations in prevalence are the norm, and prior detection of STEC O157 is not predictive of its subsequent detection.^5-8^ One model estimated that pathogen extinction within a single population occurs with 100% probability within 20-160 days.^9^ Moving beyond a single farm, studies have shown similar strains detected at neighboring farms,^6,10^ and multiple models suggest cattle movement is an important component of prevalence.^11,12^ However, modeling persistence rather than prevalence shows that even inter-farm cattle movements cannot explain STEC persistence,^12^ suggesting that if long-term local persistence occurs, it entails mechanisms not yet understood.

With the broader potential for local persistence unclear, we tested the hypothesis that LPLs are a common feature of regional STEC O157 disease dynamics. We also generalized the LPL concept for use in areas where primarily only human case surveillance data are available; e.g., the entire United States. In doing so, we determined that local persistence is more common than our current understanding of persistence mechanisms would suggest. We also identified regional variation in LPLs and laid a path for elucidating national STEC dynamics.

## Methods

The study population included all available STEC O157:H7 cases reported to the Minnesota Department of Health (MDH), designated as the “local” population of interest (Supplement). For every Minnesota case, we randomly sampled three “external” cases reported to health departments in the other 49 states. Selection was limited to cases with whole genome sequenced isolates reported to PulseNet 2010-2019 (Supplemental Data). This study was determined to be exempt by the University of Minnesota IRB.

We assembled, QCed, typed, aligned, and determined the clade of sequences as previously described,^2^ principally using the Bactopia v3.0.0 pipeline (Supplement).^13^ The resulting core SNP alignment was used in BEAST2 v2.6.7 to generate a time-calibrated phylogeny (Supplement).^14^ A sensitivity analysis was conducted using whole genomes instead of the core SNP alignment (Supplement).

We identified LPLs based on the following criteria: 1) a single lineage of the tree with a most recent common ancestor (MRCA) with ≥80% posterior probability; 2) contained ≥3 non-downsampled sequenced isolates; 3) contained >⅔sequences from the locale of interest (Minnesota in this case); 4) sequences spanned a period of ≥1 year; and 5) all isolates were ≤100 core SNPs from one another. These parameters were chosen after testing various combinations and can be modified based on specific public health goals (Supplement).

We calculated the proportion of Minnesota and external isolates that were associated with LPLs, along with Clopper-Pearson exact binomial 95% confident intervals (CIs). We identified differences between LPL and non-LPL isolates in outbreak characteristics, clinical outcomes, and genetic characteristics using chi-square tests, Fisher’s exact or analysis of variance (ANOVA) (Supplement). To determine whether LPL isolates were geographically clustered, as would be expected if they represent local patterns of disease ecology, we assessed the spatial distribution of LPL and non-LPL isolates based on the zip code of the reported case’s residence. We identified spatial clusters of LPL and non-LPL cases using Poisson spatial scan statistics (Supplement).

## Results

Of 3,039 STEC cases reported to the Minnesota Department of Health (MDH) from 2010 through 2019, we identified 373 O157:H7 isolates that had been sequenced and were available on NCBI. Of these, 10 (2.6%) could not be downloaded, failed assembly, or failed QC. STECFinder confirmed the identity of the remaining 363 as STEC O157:H7. We randomly sampled 1,245 isolates external to Minnesota that had been reported and sequenced through PulseNet by other states, of which 52 (4.2%) could not be downloaded, failed assembly, or failed QC and 29 (2.3%) were clade A (Supplement). This left 1,164 external isolates for analysis.

### Locally Persistent Lineages Were Common

We identified 15 Minnesota STEC O157:H7 LPLs (Figure 1A). These LPLs were associated with 35.3% (95% CI 30.3%, 40.4%) of Minnesota STEC O157:H7 cases reported during the study period, after reincorporating downsampled isolates (Supplemental Table S1). LPLs were also associated with 21 external isolates, of which 10 were reported in other Upper Midwest states. Including both Minnesota and external cases, LPLs ranged in size from 3 to 23 isolates, with an average of 9.9 (SD 6.8). They produced cases for 1.3 to 8.6 years (mean 3.8, SD 2.8). Clade G contained the greatest number of LPLs, with 7, accounting for 47.5% of its Minnesota cases (95% CI 38.3%, 56.8%) (Supplement, Supplemental Table S1). A sensitivity analysis with whole genomes rather than SNPs suggested uncertainty in the LPLs identified from the uncommon clades (Supplement).

**Figure 1.**
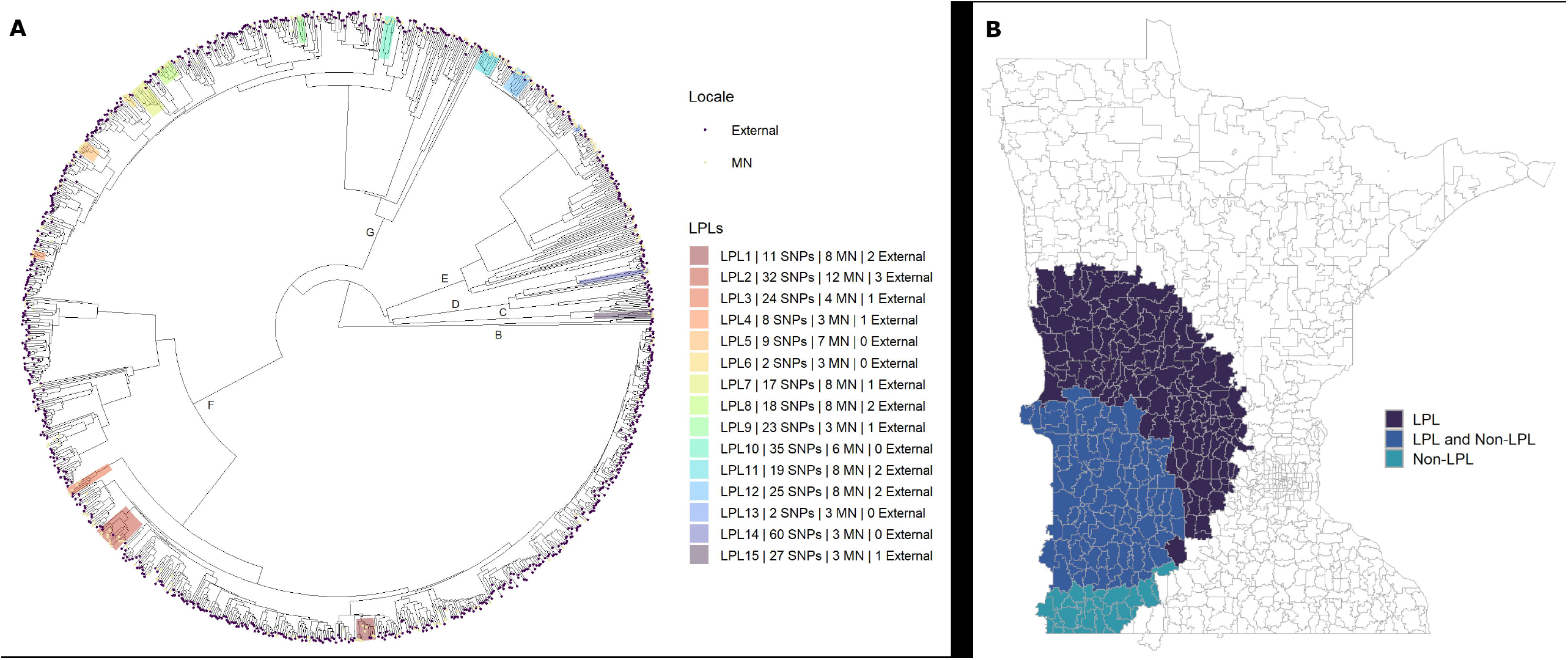
Locally persistent lineages (LPLs) identified from the O157:H7 STEC phylogeny and their spatial distribution. A) We identified 15 LPLs, which included 128 Minnesota (35.3%) and 21 external (1.8%) isolates. The number of isolates used to define the LPLs is shown in the legend; the inclusion of downsampled isolates is not shown on the figure but is included in the total. O157 clades are indicated on inner branches. LPLs occurred in all clades except clade B. Clade F contained 3 LPLs, accounting for 45 of its 151 Minnesota isolates (29.8%) and produced cases over an average of 8.1 years. Clade G contained 7 LPLs, accounting for 57 of its 120 Minnesota isolates (47.5%), and produced cases over an average of 3.0 years. Clade E contained 3 LPLs, accounting for 20 of its 78 Minnesota isolates (25.6%), and produced cases over an average of 2.5 years. Clades C and D each contained 1 LPL with 3 cases. B) Poisson-based spatial scan statistics limited to 10% of the population identified individual clusters of LPL and non-LPL isolates, which partially overlapped. The incidence of an LPL-associated O157:H7 STEC infection was 2.97 times greater (p = 0.001) within the cluster relative to the risk in the entire state. The incidence of a non-LPL associated O157:H7 STEC infection was 3.62 times greater (p = 0.001) within the cluster relative to the risk in the entire state.

### Epidemiological and Genetic Differences

A greater proportion of LPL than non-LPL isolates were linked to outbreaks, and the geographic source of the outbreaks (Minnesota or multi-state) differed by LPL status (p < 0.001) (Table 1). Forty (31.3%; 95% CI 23.4%, 40.0%) LPL isolates were linked to outbreaks with Minnesota sources, compared to only 5.1% (95% CI 2.7%, 8.7%) of non-LPL isolates. Supporting the veracity of the LPLs, there were no LPL isolates linked to multi-state (non-Minnesota) outbreaks, whereas 16.6% (95% CI 12.1%, 22.0%) of non-LPL isolates were linked to multi-state outbreaks.

**Table 1.** Comparison of LPL and non-LPL O157:H7 STEC isolates from Minnesota, 2010-2019.

|  | **LPL (n=128)** | **Non-LPL (n=235)** | **Chi-square**<br>**p-value** |
| --- | --- | --- | --- |
| **Outbreak-associated, n (%; 95% CI)** |  |  | <0.001 |
| Minnesota outbreak | 40 (31.3%; 23.4%, 40.0%) | 12 (5.1%; 2.7%, 8.7%) |  |
| Multi-state outbreak | 0 (0%; 0%, 2.8%) | 39 (16.6%; 12.1%, 22.0%) |  |
| Sporadic | 88 (68.8%; 60.0%, 76.6%) | 184 (78.3%; 72.5%, 83.4%) |  |
| **Minnesota outbreak type, n (%; 95% CI)** |  |  | 0.596 <sup>a</sup> |
| Animal contact | 13 (32.5%; 18.6%, 49.1%) | 3 (25.0%; 5.5%, 57.2%) |  |
| Foodborne | 17 (42.5%; 27.0%, 59.1%) | 4 (33.3%; 9.9%, 65.1%) |  |
| Person-to-person | 10 (25.0%; 12.7%, 41.2%) | 5 (41.7%; 15.2%, 72.3%) |  |
| **Clinical outcomes, n (%; 95% CI)** |  |  |  |
| Bloody diarrhea | 85 (66.4%; 57.5%, 74.5%) | 167 (71.1%; 64.8%, 76.8%) | 0.759 |
| Hospitalized | 53 (41.4%; 32.8%, 50.4%) | 106 (45.1%; 38.6%, 51.7%) | 0.744 |
| HUS | 24 (18.8%; 12.4%, 26.6%) | 51 (21.7%; 16.6%, 27.5%) | 0.684 |
| **stx profile, n (%; 95% CI)** |  |  | <0.001 <sup>a</sup> |
| *stx1a* | 1 (0.8%; 0%, 4.3%) | 1 (0.4%; 0%, 2.3%) |  |
| *stx1a/stx2a* | 50 (39.1%; 30.6%, 48.1%) | 31 (13.2%; 9.1%, 18.2%) |  |
| *stx1a/stx2c* | 24 (18.8%; 12.4%, 26.6%) | 50 (21.3%; 16.2%, 27.1%) |  |
| *stx2a* | 8 (6.3%; 2.7%, 11.9%) | 38 (16.2%; 11.7%, 21.5%) |  |
| *stx2c* | 2 (1.6%; 0.2%, 5.5%) | 19 (8.1%; 4.9%, 12.3%) |  |
| *stx2a/stx2c* | 38 (29.7%; 21.9%, 38.4%) | 90 (38.3%; 32.1%, 44.8%) |  |
| Other *stx1/stx2* | 5 (3.9%; 1.3%, 8.9%) | 6 (2.6%; 0.9%, 5.5%) |  |
| **Other virulence genes, n (%; 95% CI) <sup>b</sup>** |  |  |  |
| *ehxA* | 125 (97.7%; 93.3%, 99.5%) | 221 (94.0%; 90.2%, 96.7%) | 0.192 <sup>a</sup> |
| *espP* | 122 (95.3%; 90.1%, 98.3%) | 227 (96.6%; 93.4%, 98.5%) | 0.575 <sup>a</sup> |
| **ARGs present, n (%; 95% CI)** | 6 (4.7%; 1.7%, 9.9%) | 42 (17.9%; 13.2%, 23.4%) | 0.001 |
| **Stress tolerance genes, n (%; 95% CI) <sup>c</sup>** |  |  |  |
| *emrE* | 110 (85.9%; 78.7%, 91.4%) | 191 (81.3%; 75.7%, 86.1%) | 0.326 |
| *qacE* | 0 (0%; 0%, 2.8%) | 18 (7.7%; 4.6%, 11.8%) | 0.003 |
| Mercury tolerance <sup>d</sup> | 4 (3.1%; 0.9%, 7.8%) | 19 (8.1%; 4.9%, 12.3%) | 0.104 |

Abbreviations: ARG, antimicrobial resistance gene; CI, confidence interval; HUS, hemolytic uremic syndrome; LPL, locally persistent lineage
<sup>a</sup> Tested using Fisher's exact test.
<sup>b</sup> Virulence genes *eae*, *iha*, and *tir* were present in all isolates.
<sup>c</sup> Tolerance genes for arsenic (*arsC* and *arsR*) and tellurium (*terD*, *terW*, and *terZ*) were present in all isolates. Tolerance genes for acid (*asr*) were present in 99.7% of isolates.
<sup>d</sup> Includes any combination of genes *merA*, *merB*, *merC*, *merD*, *merE*, *merP*, *merR*, and *merT*.

LPL and non-LPL isolates also differed genetically (Table 1). There was a substantial difference (p < 0.001) in Shiga toxin gene (stx) profile, a key virulence determinant, with 39.1% (95% CI 30.6%, 48.1%) of LPL isolates carrying *stx1a/stx2a* combinations, compared to 13.2% (95% CI 9.1%, 18.2%) of non-LPL isolates. Despite the differences in *stx*virulence profiles, there were no differences in prevalence of hemolytic uremic syndrome, hospitalization, or bloody diarrhea between cases infected with LPL vs. non-LPL STEC. LPL isolates were also less likely than non-LPL isolates to carry any antimicrobial resistance genes (ARGs) and the *qacE* gene.

### Spatial Distribution of LPL and Non-LPL STEC Overlapped

We identified a cluster of LPL cases in the western part of the state where incidence within the cluster was 2.97 times (p = 0.001) the incidence in all of Minnesota (Figure 1B). We also found a non-LPL cluster in the southwest part of the state where incidence was 3.62 times (p = 0.001) that in the entire state. The two clusters partially overlapped. However, the most likely cluster identified in a direct comparison of LPL to non-LPL cases was not statistically significant (RR 2.00, p = 0.081). All results were consistent in sensitivity analyses of different window sizes.

## Discussion

In this study, we identified 15 STEC O157:H7 LPLs in Minnesota, supporting our initial hypothesis that LPLs are a common feature of STEC disease systems. LPL-associated infections were epidemiologically important, accounting for 35.3% of all Minnesota cases and 44.0% of cases linked to identified outbreaks. Several aspects of our findings support LPLs as genuine ecological phenomena linked to local reservoirs.

First, we observed LPLs that were associated with cases up to 9 years apart, consistent with results from Alberta where LPLs persisted up to 13 years.^2^ Long-term persistence within individual hosts and herds has been ruled out; thus, our results suggest ecosystem-level persistence. STEC colonization in cattle persists in most animals for <60 days.^4^ Longitudinal studies report large fluctuations in STEC prevalence, ^5,7,8^ and models of the role of the environment in maintaining STEC on farms predict fade-out in all cases.^9^ Even accounting for the movement of cattle between farms, STEC persistence cannot be explained by cattle alone.^12^The identification of LPLs is a starting point for investigating the factors that enable STEC persistence within ecosystems.

Second, while our LPL criteria required that more than two-thirds of isolates in a lineage had been reported in Minnesota, thus allowing up to 32% to be external, external isolates accounted for only 14% of LPL isolates. This demonstrates even greater dominance by Minnesota isolates than expected. Additionally, almost half of external LPL cases were reported in neighboring states in the Upper Midwest, potentially due to visitors to Minnesota from neighboring states or ecosystem-level reservoirs that cross state lines.

Third, LPLs contained no isolates linked to outbreaks with non-Minnesota sources, which if they had, would have suggested erroneous LPL identification. Most cases linked to outbreaks with Minnesota sources, 76.9%, were associated with LPLs. This emphasizes the public health importance of these strains. LPLs are location-specific targets defined based on repeated human infections. If their specific reservoir populations can be identified, control measures could potentially be implemented to eradicate them, or animals in those populations could be screened prior to slaughter. Widespread adoption of whole genome sequencing also means that public health authorities could identify cases associated with LPLs and direct their investigations to local sources, which could be particularly advantageous when resources for public health are limited.

Fourth, both LPL and non-LPL spatial clusters were in areas of the state with high densities of domestic ruminants, and the cluster of LPL-associated cases overlapped an area with a high concentration of dairy cattle (i.e., Minnesota’s “dairy belt”). Further investigation is needed to determine whether LPLs are linked to dairy or other agricultural operations, but this clustering suggests an association between LPLs and a putative local reservoir. Additionally, the spatial clustering we found is much broader than a single farm, implying ecosystem-level persistence.

Finally, we identified important genetic differences between LPL and non-LPL isolates, suggesting common selective pressures among LPLs. The stx profile of LPL vs. non-LPL isolates differed significantly, and ARG and *qacE* prevalence were less common among LPL than non-LPL isolates. These factors could affect survival in the ruminant reservoir or environment.^15^ Additionally, clade G isolates were more likely to be associated with LPLs than isolates in other clades, which is consistent with results from Alberta.^2^

This study was limited by a lack of data from putative animal and environmental reservoirs. Such reservoir data is not widely available, as evidenced by the availability of only 4 animal or environmental isolates from Minnesota. This was the motivation for developing a method that relied only on human surveillance data, which is common in the U.S. We were also limited in available phylogenetic methods. With the gradual introduction of whole genome sequencing in surveillance, sequenced isolates were skewed toward the latter part of our study period, precluding the use of phylogeographic methods. The combination of the size of STEC genomes and the number of isolates sequenced also prevented us from using whole genomes with integrated BEAST2 phylogenetic models. Our sensitivity analysis with whole genomes and step-wise analysis suggested uncertainty in the uncommon clades. Any public health action based on LPLs should use additional epidemiological information to verify LPLs in uncommon clades.

This study fills a gap in our knowledge of STEC persistence. We have shown that STEC strains can persist in a limited geographic area for many years and that such persistent strains contribute significantly to local disease burden. From differences between Minnesota and our previous study in Alberta, we have learned that the role of persistent strains in local epidemiology is location dependent. Minnesota has more typical STEC incidence and cattle population, relative to the rest of the U.S., increasing the generalizability of our results here. Despite using locality to define them, the impact of LPLs is national, as bacteria local to one area can contaminate food items that are shipped throughout the country. Thus, controlling LPLs where they are local has the potential to prevent disease on a much larger scale.

## Acknowledgements

We thank Phillip I. Tarr, Craig Hedberg, Jeffrey Bender, and Noelle Noyes for their contributions to the study and feedback on the manuscript. We gratefully acknowledge the PulseNet participating laboratories, whose data was used for the creation of this publication. The authors acknowledge the Minnesota Supercomputing Institute (MSI) at the University of Minnesota for providing resources that contributed to the research reported in this paper.

## Data Availability

All sequencing data are available on NCBI at the run IDs listed in Supplemental Data. Metadata must be requested from the CDC PulseNet team.

