## Supplement for "The conundrum of Shiga toxin-producing *Escherichia coli* O157:H7 persistence: Evidence for locally persistent lineages"

### Study Population and Case Definition

We designated cases reported in Minnesota as “local” and cases reported in the other 49 states as “external”, though the utility was designed to allow any state or region to be designated as the locality of interest. We included all available Minnesota O157 STEC sequences and randomly sampled external sequences at a ratio of approximately 3:1. Initial sampling was based on the serotype identified in the PulseNet database. The study population included O157 STEC cases reported to the Minnesota Department of Health (MDH) for Minnesota cases, or health departments outside of Minnesota for external cases. MDH and other health departments subsequently reported cases to the Centers for Disease Control and Prevention (CDC) PulseNet surveillance system.^1^ MDH obtains extensive metadata about reported cases, including home zip code and whether a case was linked to a known outbreak.

Cases with an isolation date in 2010 through 2019 were included. Only cases reported within the 50 U.S. states with isolates that had been whole genome sequenced were included. Minnesota isolates were slightly overrepresented in 2010 and underrepresented during 2011-2016 (Supplemental Figure S1). Twelve percent of external isolates (n=141) were reported from neighboring Upper Midwest states (Iowa, South Dakota, North Dakota, Wisconsin).

Cases were defined as individuals meeting the Council of State and Territorial Epidemiologists (CSTE) confirmed case definition that was in use at the time the case was reported; CSTE STEC case definitions applying to this study were adopted in 2006, 2014, and 2018.^2–5^ In most instances, cases sought care and provided a fecal sample, which was tested by their clinician. Fecal samples, and in some cases bacterial isolates, were delivered to public health laboratories, where, if necessary, fecal samples were cultured to obtain an isolate, and all isolates were whole genome sequenced per PulseNet procedures.^6^ Public health laboratories participating in PulseNet adopted routine STEC sequencing between 2016 and 2019; sequences from isolates collected prior to routine sequencing were usually retrospectively sequenced due to particular interest in the case, such as involvement in an outbreak, severe outcome, or unique source of infection. MDH began routine STEC sequencing in 2016 and sequenced an additional 39 isolates from 2010 to 2015 to provide more data for the early years of the study. This study was deemed exempt by the UMN IRB.

### Bioinformatics Processing

#### Sequence Assembly, Validation, and Characterization

SRA Toolkit v3.0.0 was used to download all sequences from NCBI. We used Bactopia v3.0.0^7^ for assembly and quality checking. First, FastQC v0.12.1 evaluated raw reads based on read count, sequence coverage, and sequence depth. Failed reads were excluded from subsequent assembly. We *de novo* assembled raw reads with Shovill v1.1.0 within Bactopia, which included use of Trimmomatic to trim adapters and read ends with quality <6. Reads with overall quality scores <10 after trimming were discarded. After Bactopia assembly, we assessed assembly quality based on number of contigs (≤400), genome size (≥5.0 Mb), N50 (>30,000), and L50 (≤50). Low-quality assemblies were excluded.

STECFinder v1.1.025 confirmed serotype using the *wzx* or *wzy* O-antigen genes and detection of the H-antigen.^8^ For the detection of virulence genes, antimicrobial resistance genes (ARGs), and stress tolerance genes, we used AMRFinder Plus v3.12.8 within the Bactopia pipeline,^9^ using *Escherichia coli* O157:H7 str. Sakai (NC_002695.2)^10^ as the reference genome and limiting results to matches with >90% sequence coverage and identity. ARGs that did not correspond to a specific antibiotic, including efflux pump genes and *blaEC* were excluded from the analysis.

#### Sequence Alignment

We generated a core SNP alignment with recombinant blocks removed using Bactopia’s Snippy workflow, which incorporates Snippy v4.6.0, Gubbins v3.3.0, and IQTree v2.2.2.7.^7,11^ We reduced the final alignment to core genome SNPs using SNP-Sites v2.5.1 The number of core SNPs between isolates was calculated using PairSNP v0.3.1. Across all 1,527 isolates, we identified 11,437 core genome SNPs.

Prior to phylogenetic analysis, we pruned alignments of similar sequences agnostic to location in a process called “downsampling” (Supplemental Figure S2).^12^ The downsampling process we developed identified sequences within 2 SNPs of another sequence. Additionally, downsampled sequences were from cases reported <60 days after a matching sequence, with no more than 12 months of continuous cases from the initial matching sequence. We noted the matching sequence retained within the alignment to later reincorporate the downsampled sequences into the statistical analysis of LPLs. If any isolate within a downsampled group was linked to a known outbreak, all isolates in the downsampled group were considered to have been part of the same outbreak. We downsampled 116 Minnesota (32.0%) and 354 external (30.4%) isolates due to high sequence similarity.

### Phylogenetic Analysis

We used BEAST2 v2.6.7 to generate a time-calibrated phylogeny.^13^ To allow for changes in population dynamics over time, we used a Bayesian coalescent skyline tree model with 3 skyline pieces. For the substitution model, we used an HKY prior with empirical frequencies and discrete Gamma site model with four categories. For the clock model we used a relaxed log-normal prior with initial value 1.5×10^–5^ and log-normal distribution (M=1.5×10^–5^, S=1.5, with mean in real space).^12^ Four chains were run for a total of 162.8 million iterations with 10% burn-in and sampling every 40,000 steps. The chains were run till convergence with >200 estimated sample size for all non-skyline parameters.

The root of the time-calibrated phylogenetic tree was estimated to be 1846 (95% HPD 1811, 1883). The estimated clock rate for the core genome was 8.00x10^-5^ (95% HPD 7.16x10^-5^, 8.96x10^-5^) substitutions/site/year.

### Locally Persistent Lineage (LPL) Identification

A maximum clade credibility (MCC) tree was generated from the combined chains in BEAST2, which was used for the definition of the LPLs. We built a custom utility within R to identify LPLs, which were highlighted on phylogenies using the treeio and ggtree packages in R.^14–16^ The utility is generalizable beyond this study, allowing specification of all detection criteria and the locale of interest. The locale could be another state, a region (e.g. the Upper Midwest), or an entire country if interested in international comparisons. The selection of isolates for analysis would simply need to support the chosen location. We randomly selected isolates from outside of Minnesota for comparison, limiting our selections to other human-derived isolates identified through the same reporting mechanism, i.e. PulseNet. Having local and external isolates from the same source reduces opportunities for sampling or selection bias.

### LPL Characteristics

#### LPL Incidence Rate in Minnesota vs. Alberta

We calculated the incidence rates of LPL- and non-LPL-associated O157 STEC cases in Minnesota and compared those to the corresponding incidence rates from Alberta, Canada.^12^ We calculated the incidence rates from 2010-2019 by multiplying the proportion of Minnesota cases associated with an LPL, or 1 minus that proportion for the non-LPL rate, by the overall STEC incidence rate (7.08 per 100,000^17^), multiplied by the proportion of reported STEC cases in the state that were infected with STEC O157:H7 (40.2%^18^). We compared these incidence rates to similarly calculated rates for Alberta, Canada, where LPLs have previously been investigated: LPL proportion (74.7%^12^) times overall STEC incidence (8.45 per 100,000^19^) times the proportion of cases that were infected with STEC O157:H7 (40.3%^20^).

For Minnesota, we estimated incidence of 1.0 LPL-associated and 1.8 non-LPL-associated STEC O157:H7 cases per 100,000. In Alberta, these values were 2.54 and 0.86 per 100,000, respectively. Thus, the difference in the proportion of O157:H7 STEC cases associated with LPLs, 35.3% in Minnesota and 74.7% in Alberta, is indicative of true differences in LPL incidence. This was not unexpected given the importance of cattle as a reservoir. Alberta has over twice the number of cattle, with 4.5 million head in 2020,^21^ as Minnesota, with 2.1 million head in 2024,^22^ and it is well-established that living in a region with a high density of ruminants increases the risk of STEC.^18,23,24^ LPLs could help explain the mechanism for this increased risk. Large ruminant populations are more likely than small populations to exceed the critical community size required for pathogen maintenance, resulting in less stochastic fade-out.^25,26^ Strain persistence in ruminant-dense regions would provide greater opportunities for both direct and indirect human infections. However, a systematic study is needed to compare LPLs across geographic areas to identify regional factors that influence LPL-associated disease burden.

#### LPL Differences Between Clades

O157 STEC clades were assigned based on SNPs identified by Strachan et al.^27^ Based on this scheme, clades are lettered A through G, with subclades indicated by roman numerals; e.g. G(vi). To avoid inferring deeper evolutionary histories than our data could support, we excluded isolates in clade A (n=29), which included only external isolates. After QC, clade B also contained only external isolates; thus, no LPLs were identified in clade B. We compared the proportion of cases associated with LPLs across clades using a chi-square test. For each LPL, we determined the number of isolates per LPL and duration from earliest isolate to latest isolate. For these measures, we calculated the mean value and standard deviation (SD) for all O157 STEC and for each clade. We compared the means between clades using analysis of variance (ANOVA), limiting the analysis to clades with >1 LPL. If the Shapiro-Wilk test indicated non-normality in ANOVA residuals, we investigated the source (e.g. an outlier) and performed a sensitivity analysis (e.g. without the outlier).

Clade F was the most common clade for both external and Minnesota isolates. The proportion of Minnesota isolates identified as clade E was three times that of external isolates (Supplemental Figure S3). Clade G contained the greatest number of LPLs, with 7, followed by E and F, with 3 LPLs each. Clade G also had the greatest proportion of its Minnesota cases associated with LPLs (47.5%; 95% CI 38.3%, 56.8%) (Supplemental Table S1). In Clade F, 29.8% (95% CI 22.6%, 37.8%) of Minnesota isolates were associated with LPLs, as were 25.6% (95% CI 16.4%, 36.8%) of Minnesota isolates in clade E. Clades C and D each contained 1 LPL with 3 Minnesota isolates each. The proportion of Minnesota isolates associated with LPLs differed significantly by clade (p = 0.006). Clade G’s high number of LPLs was consistent with our findings in Alberta, where >90% of STEC O157:H7 isolates, including all LPL isolates, belonged to clade G.^12^ Future work is needed to determine whether there are genetic factors within this clade that predispose its isolates to developing LPLs.

Clade F LPLs differed from those in other clades (Supplemental Table S1). They had more isolates per LPL (mean 17.7, SD 5.9) and produced cases for more years (mean 8.1, SD 0.8); only the ANOVA comparing differences in mean duration across clades was statistically significant (p = 0.008). It is unclear whether there are genetic factors influencing longer duration of persistence in clade F or whether this is chance based on the reservoirs where clade F LPLs are established.

#### Genetic Differences Between LPLs

We calculated the frequency of LPL vs. non-LPL isolates that carried each combination of *stx* genes, other virulence genes, ARGs, and stress tolerance genes. For each proportion, we calculated an exact binomial 95% CI and used a chi-square test for independence or Fisher’s exact test to compare proportions between LPL and non-LPL groups.

The *stx1a/stx2a* profile was also significantly more common among LPL isolates, and isolates with only an *stx2* gene were more commonly not associated with LPLs. Virulence is attenuated in isolates with both *stx1* and *stx2* genes relative to only *stx2*,^28–30^ though we found no difference in clinical outcomes between LPL- and non-LPL-associated cases. The *stx1/stx2* profile of LPL isolates may indicate that high virulence is not an important trait for establishing or maintaining STEC populations in the ecosystem. For example, *stx2*-carrying isolates were more common among cattle than those carrying only *stx1* or a combination.^31^ However, our work in Alberta found that LPLs adopted more virulent *stx* profiles over time,^12^ so it is also possible that selection on virulence is niche-specific. Our examination of other virulence genes in the current study identified no differences between LPL and non-LPL isolates.

We found differences in genes for antimicrobial resistance and quaternary ammonium compound (QAC) tolerance. In both cases, non-LPL isolates had significantly more resistance/tolerance genes. Antimicrobial resistance has been associated with persistence within cattle,^32^ and while persistence within the animal host is a complex trait,^33,34^ the reverse association between resistance and LPLs suggests that the persistence of LPLs is not driven by intra-host persistence, or if it is, other virulence factors are responsible. QAC tolerance in *E. coli* is a recognized issue and is often coupled with antimicrobial resistance.^35^ However, *qacE* does not necessarily confer QAC resistance.^36^ Its potential role in LPL persistence, if any, is thus unclear.

#### LPL Spatial Distribution

Based on the zip code tabulation area (ZCTA) of the reported case’s residence, we created a bivariate choropleth map using the biscale package (Supplemental Figure S4).^37^ We used Poisson spatial scan statistics in the SpatialEpi package^38^ to identify separate spatial clusters of LPL and non-LPL isolates. We used the population within each ZCTA as the reference population and limited clusters to containing up to 10% of the population. Sensitivity analyses adjusted this limit to 20% and 50% of the population. The standardized incidence ratio (SIR) of each spatial cluster gave the ratio of the LPL or non-LPL incidence rate within the cluster relative to the state average. We also directly compared LPL to non-LPL isolates using Bernoulli scan statistics in the smacpod package,^39^ which identified spatial clusters in which the ratio of LPL to non-LPL isolates was statistically greater inside the cluster than outside the cluster, given as a relative risk (RR). The maximum cluster size was set at a radius of 50 miles; 25 and 100 mile radii were tested during sensitivity analysis. The alpha level for the primary cluster in all analyses was set at 0.05.

### Sensitivity Analyses

#### LPL Identification Parameters

Our interest in defining the LPLs was to identify strains that are persisting within Minnesota and contributing to the O157:H7 STEC burden, with the intent that these could be monitored to enable better public health control. The LPL definition was determined to: 1) detect persistence that goes beyond a point source outbreak (duration); 2) identify both highly active strains and strains that may just be emerging as public health threats in the state (number of isolates); and 3) limit detection to LPLs principally in Minnesota, while allowing for exportation of cases to other states (proportion local). To accomplish these goals, we explored all parameters of the LPL definition (Supplemental Table S2). We used two investigated outbreaks to validate the parameter selection, one with a known Minnesota source and one with a known source outside of Minnesota.

The results were reasonably stable across parameter ranges; only four parameter combinations produced different results. Reducing the proportion of isolates that are from the locale of interest caused a large jump in the number of LPLs and cases. The veracity of the additional LPLs was suspect, as evidenced by the inclusion of a multi-state outbreak. An option requiring a minimum of 5 tips per lineage rather than 3 caused the greatest reduction in LPLs while correctly classifying the outbreaks. We opted against using that much more stringent definition, because we are interested in monitoring strains that may be emerging. Two parameter combinations eliminated LPL-14 in clade D. Isolates within that lineage were separated by 60-62 SNPs, and the posterior support for the MRCA was 80.4%. We opted to include this LPL, as the total set of parameter combinations more frequently included it, and its exclusion would not have meaningfully altered our conclusions.

#### Comparison to Whole Genome Tree

With a sequence length of 5.6 Mb, using whole genome sequences in the BEAST2 analysis was not feasible, and so we used only core SNPs to generate the trees. To test the sensitivity of our analysis to the absence of invariant sites, we regenerated the whole genome masked alignment from Gubbins^40^ using 1000 bootstraps. The final Gubbins tree was generated by RAxML^41^ using an HKY substitution model, the same basic model we used in BEAST2. This did not incorporate a tree dating step, as the LPLs are determined from the date of isolation rather than inferred ancestral node dates.

When identifying LPLs, we used the same criteria for inclusion, replacing the node posterior value ≥80% with a bootstrap value ≥80%. The whole genome tree produced 20 LPLs (Supplemental Figure S5), which included 144 (39.7%; 95% CI 34.6%, 44.9%) of Minnesota O157 STEC cases. The LPLs also included 36 (3.1%; 95% CI 2.2%, 4.3%) external cases. The 95% CI for Minnesota cases overlaps the point value (35.3%) obtained from the SNP-based tree, and the confidence intervals for external cases overlap one another but not the point values. As with the SNP-based tree, LPLs identified from the whole genome tree were either sporadic or associated with Minnesota-linked outbreaks; none were associated with multi-state outbreaks.

Of the 149 isolates associated with an LPL from the SNP-based tree, 11 (7.4%), including 9 Minnesota cases, were not associated with an LPL from the whole genome tree. This included the entirety of LPLs 14 and 15 in clades D and C, respectively, and outer isolates in LPLs 2, 3, and 10. All had the same clustering on both trees, but on the whole genome tree, inclusion of these isolates in the LPLs would have exceeded the 100 SNP threshold.

There were 32 cases (23 Minnesota) identified as part of LPLs on the whole genome tree that were not part of LPLs on the SNP-based tree. The greatest additions were in clade E, where 3 LPLs with 24 cases were identified from the SNP-based tree, compared to 7 LPLs with 46 cases from the whole genome tree, almost doubling the number of cases. Across all clades, 12 added cases (8 Minnesota) appeared to be due to topological changes, whereas the other 21 (16 Minnesota) were similarly structured on the SNP-based tree but had low posterior but adequate bootstrap support for their branches.

In bacterial phylodynamics, there is a trade-off between integrated analysis, such as that offered by BEAST2, and computational tractability, particularly at the scale at which we are working.^42^ Our SNP-based tree was very similar to the whole genome tree obtained from Gubbins. The differences were minimal in clades F and G, which accounted for 86% of our sequences. Clades C and D lost their LPLs entirely, and there was more restructuring in clade E than clades F or G. This suggests that our results may be less stable for clades that are uncommon in the U.S., and additional information should be used to assess their veracity. For example, one of the new clade E LPLs identified by the whole genome tree that did not have adequate posterior support on the BEAST2 tree included cases from a Minnesota-linked foodborne outbreak, suggesting that this was a true LPL missed by our SNP-based analysis. Data from additional years may also help resolve the instability but would increase computational demands, emphasizing the importance of step-wise solutions in future LPL work.

#### Animal and Environmental Isolates

Ideally, our LPLs would be supported by the presence of animal or environmental isolates from Minnesota. Unfortunately, we could identify only 5 sequences of animal or environmental origin collected during 2010-2019 in MDH’s collection. We identified 115 STEC O157:H7 animal or environmental isolates from the other 49 states that had been reported to PulseNet during the study period. All sequences were available on NCBI. Four and 111 sequences, respectively, were successfully assembled and passed QC.

Animal and environmental isolates were examined by rerunning Snippy and Gubbins with all human, animal, and environmental isolates. We created an IQTree to examine the relationships between isolates and visualized the tree in Microreact v282 with metadata for LPL, source, and geographic location. None of the animal or environment sequences clustered with LPLs (Supplemental Figure S6), neither affirming or refuting the legitimacy of the LPLs, given the small number of Minnesota isolates (n=4).

### Supplemental Figures

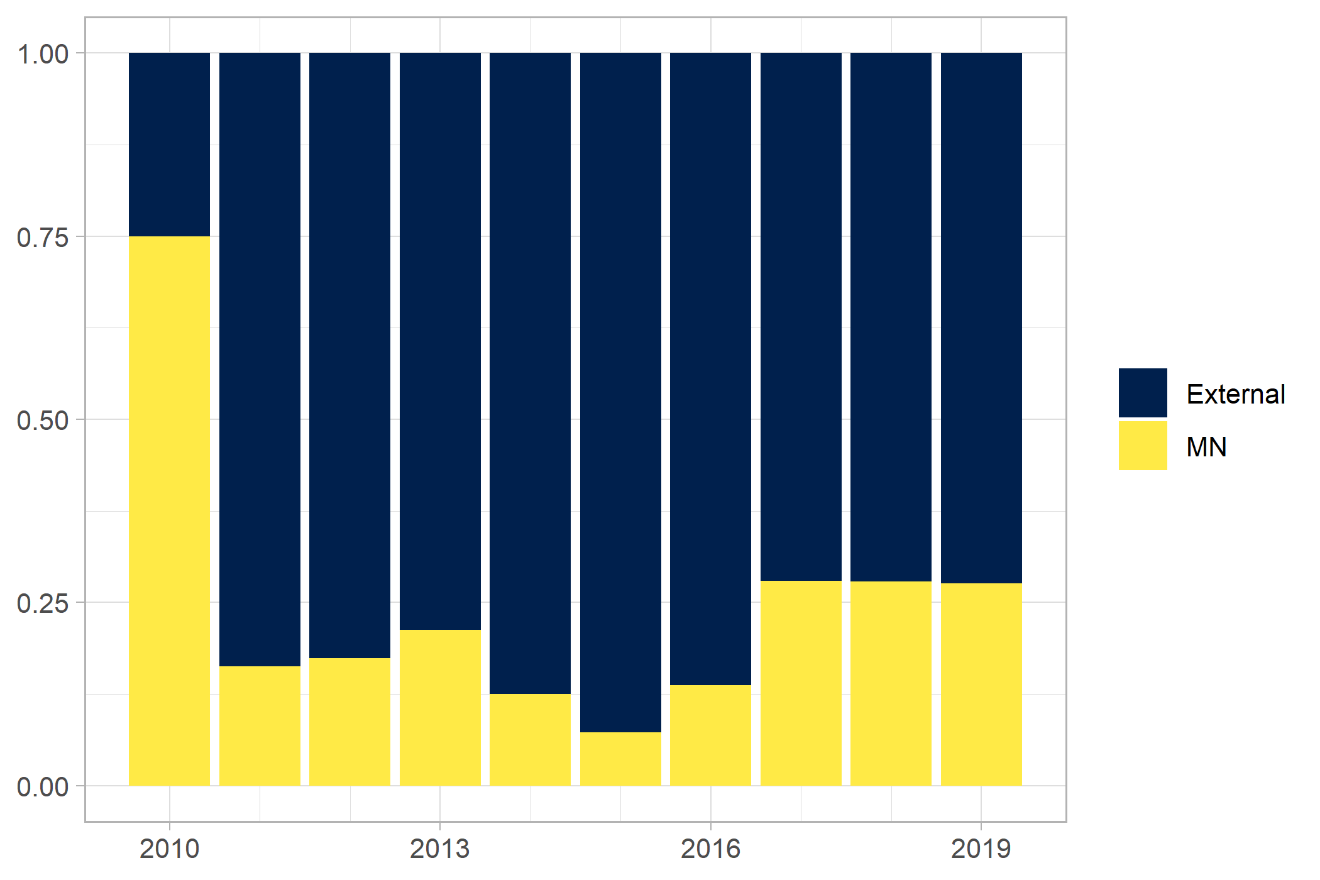

**Suppl. Fig. S1. Proportion of Minnesota vs. external sequences by year.** Overall sampling was conducted in an approximately 3:1 ratio of external to Minnesota cases. Minnesota cases accounted for 75% of isolates in 2010, 7-21% during 2011-2016, and 28% during 2017-2019.

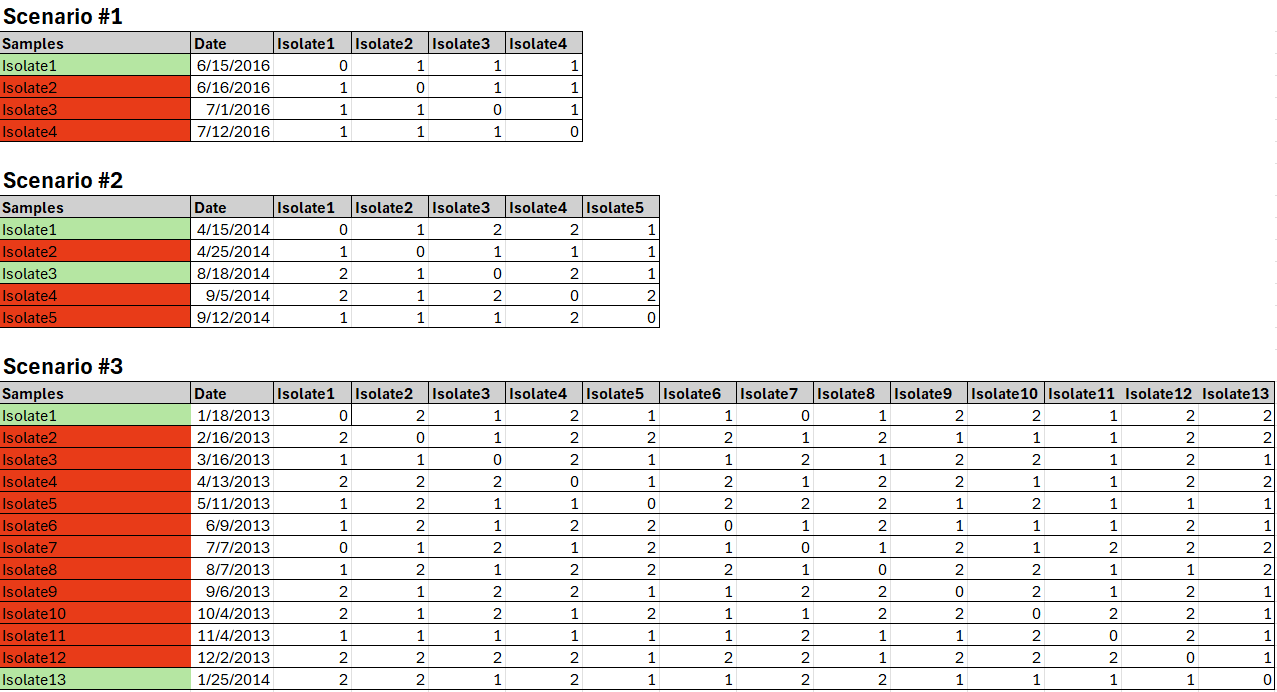

**Suppl. Fig. S2. Examples of downsampling.** Isolates were downsampled (highlighted in red) when they were within 2 SNPs and <60 days of a matching isolate (highlighted in green). If related isolates spanned more than a year, a new isolate was retained in the analysis after 12 months had elapsed since the first matching isolate (Scenario #3).

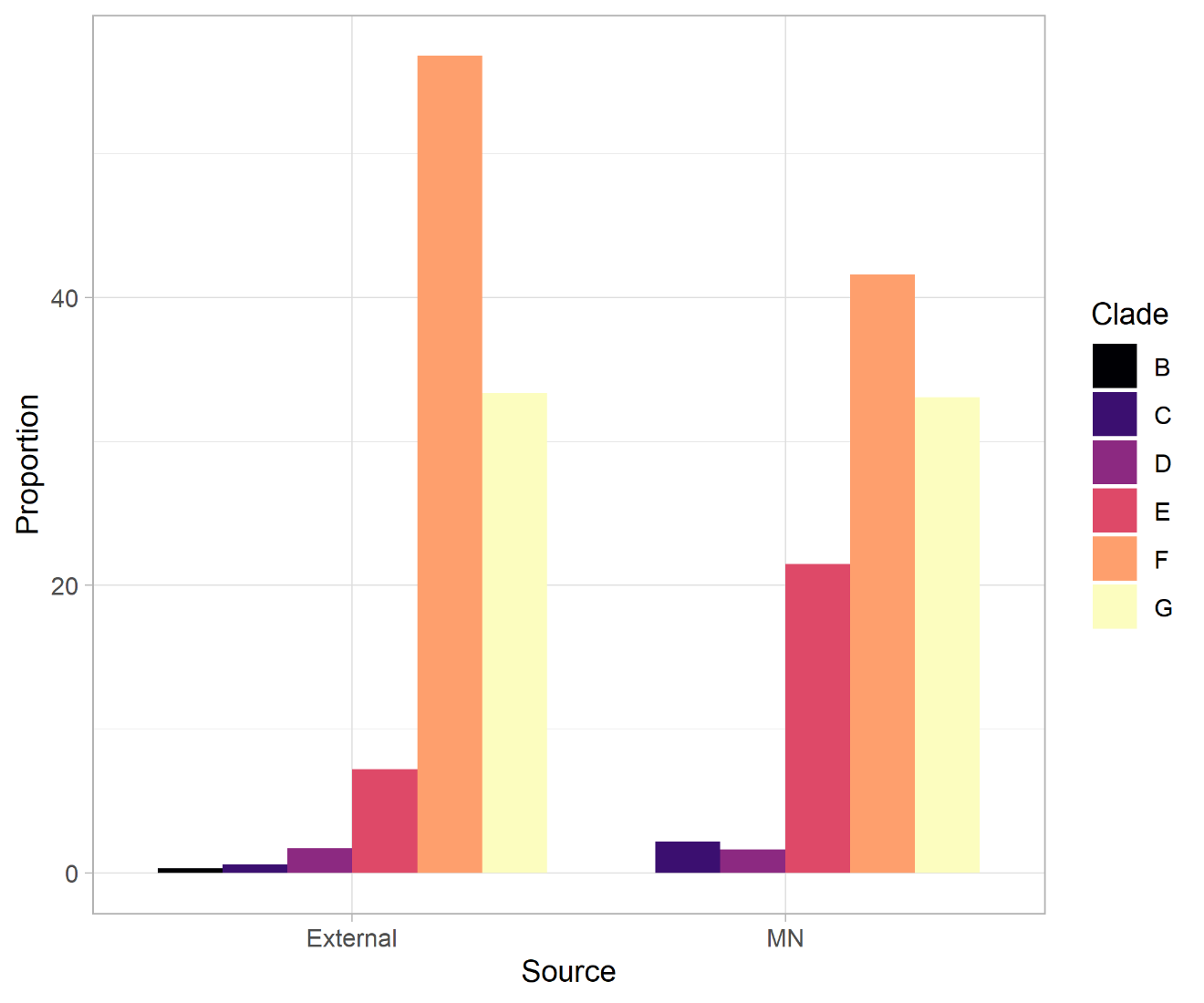

**Suppl. Fig. S3. Proportion of Minnesota vs. external sequences per clade.** F was the most common clade in both clusters, with 56.8% of external isolates and 41.6% of MN isolates. There were no clade B isolates found in MN during the study period, and the proportion of MN isolates identified as clade E (21.5%) was greater than that identified for external isolates (7.2%).

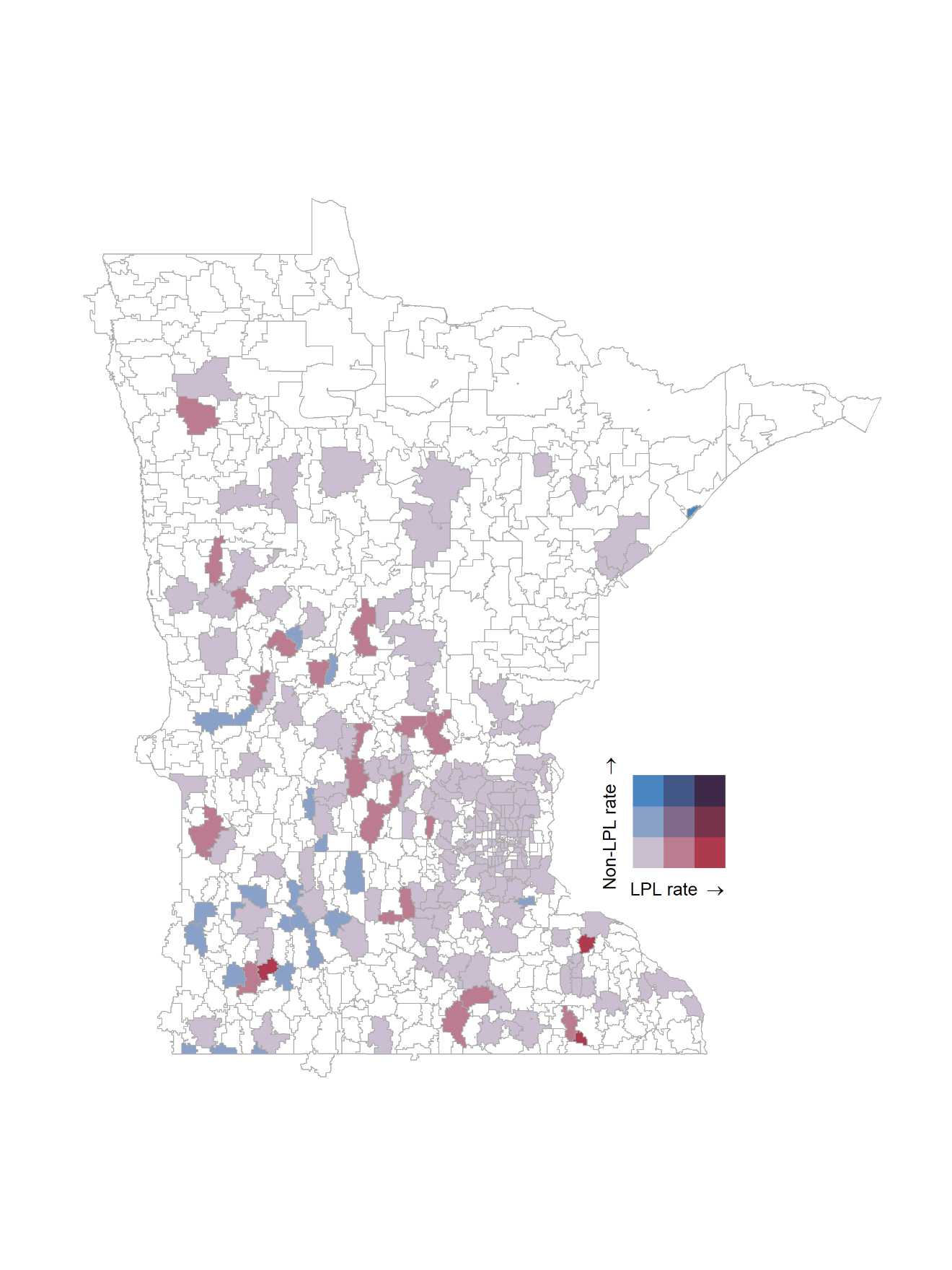

**Suppl. Fig. S4. Spatial distribution of LPL and non-LPL isolates.** Cases were aggregated to zip code tabulation areas (ZCTAs) based on home address. The incidence rate of LPL and non-LPL isolates appeared to differ in some parts of the state and overlap in others. Many ZCTAs had no STEC cases during the 10-year study period, shown in white.

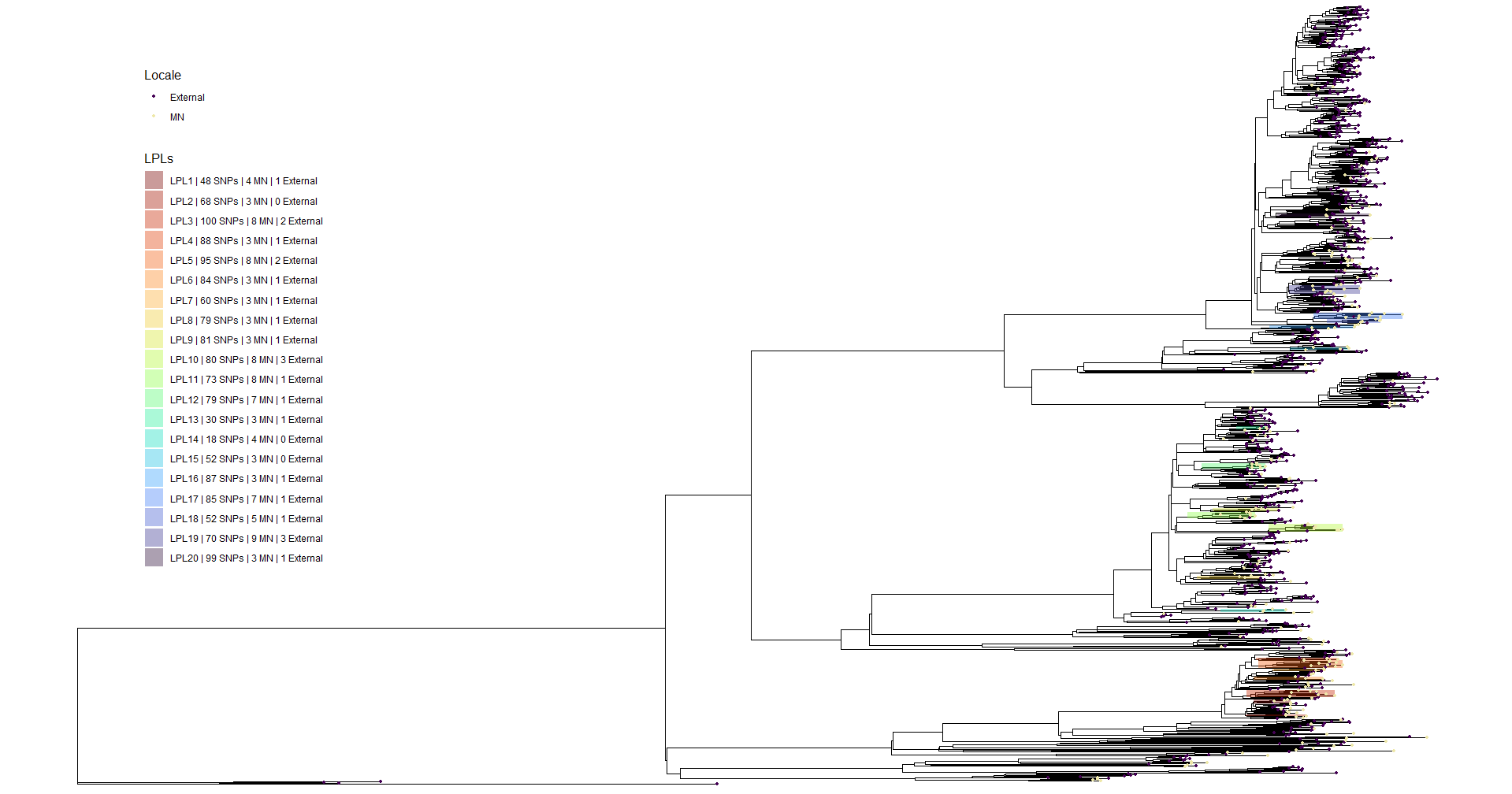

**Suppl. Fig. S5. Whole genome phylogeny with LPLs labeled.** Twenty LPLs were identified from the whole genome tree, accounting for 39.9% (95% CI 34.9%, 45.2%) of Minnesota O157 STEC cases and 3.1% (95% CI 2.2%, 4.3%) of external cases. Most notably, the LPLs in clades C and D were not identified in this tree, and 4 additional LPLs were identified in clade E.

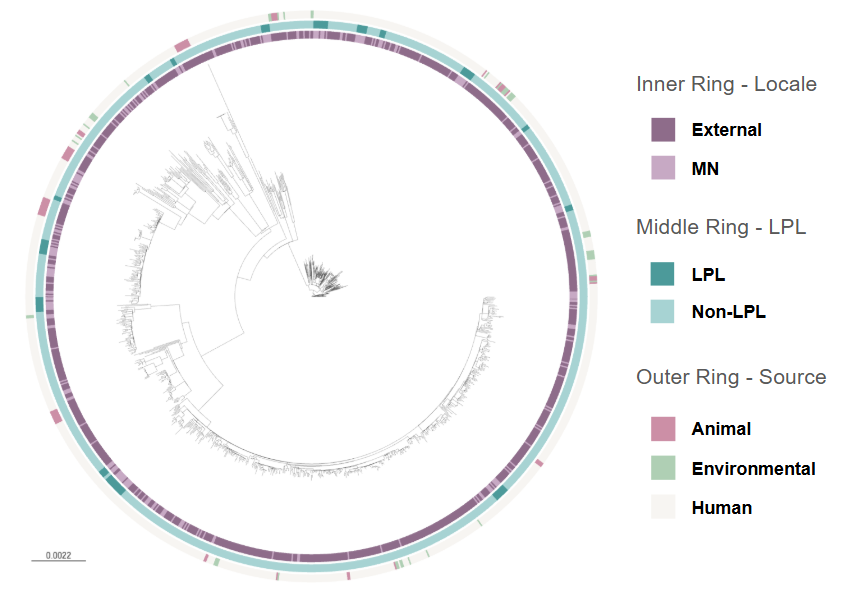

**Suppl. Fig. S6. Maximum likelihood phylogenetic tree incorporating STEC O157:H7 isolates from humans, animals, and the environment.** None of the 111 external animal and environmental isolates clustered with a Minnesota LPL. The four Minnesota animal and environmental isolates also did not cluster with any of the Minnesota LPLs.

### Supplemental Tables

**Suppl. Table S1.** Characteristics of analyzed isolates and Minnesota LPLs, by O157:H7 STEC clades

| Characteristic | All O157 | B | C | D | E | F | G | Difference by clade, p-value |
| --- | --- | --- | --- | --- | --- | --- | --- | --- |
| *Isolates* | | | | | | | |  |
| MN isolates | 363 | 0 | 8 | 6 | 78 | 151 | 120 |  |
| External isolates | 1,169 | 4 | 7 | 20 | 84 | 661 | 338 |  |
| *LPLs* | | | | | | | |  |
| LPLs, n | 15 | 0 | 1 | 1 | 3 | 3 | 7 |  |
| MN isolates, n (%; 95% CI) | 128 (35.3%;  30.3%, 40.4%) | - | 3 (37.5%;  8.5%, 75.5%) | 3 (50.0%; 11.8%, 88.2%) | 20 (25.6%; 16.4%, 36.8%) | 45 (29.8%; 22.6%, 37.8%) | 57 (47.5%; 38.3%, 56.8%) | 0.006 ^a^ |
| External isolates, n (%; 95% CI) | 21 (1.8%; 1.1%, 2.7%) | - | 1 (14.3%;  0.4%, 57.9%) | 0 (0%;  0%, 16.8%) | 4 (4.8%;  1.3%, 11.7%) | 8 (1.2%;  0.5%, 2.4%) | 8 (2.1%;  0.9%, 4.0%) | 0.038 ^a^ |
| Isolates per LPL, mean (SD) ^b^ | 9.9 (6.8) | - | 4 | 3 | 8.0 (4.4) | 17.7 (5.9) | 9.3 (6.6) | 0.137 ^c^ |
| Duration (years), mean (SD) ^b^ | 3.8 (2.8) | - | 2.1 | 2.23 | 2.5 (0.8) | 8.1 (0.8) | 3.0 (2.5) | 0.008 ^c,d^ |

^a^ Tested using Fisher’s exact test; clade B excluded.

^b^ Clades C and D each have only 1 LPL, so value for that LPL is shown rather than a distribution.

^c^ Tested using ANOVA; clades B, C, and D excluded.

^d^ Residuals were non-normal due to a high outlier in clade G; sensitivity analysis of the difference between clades E, F, and G after removal of the outlier yielded p < 0.001.

**Suppl. Table S2.** Locally persistent lineage (LPL) definition parameter testing

| Parameter | Value ^a^ | Number of LPLs | Number of Minnesota cases | Includes known Minnesota outbreaks ^b^ | Includes known multi-state outbreaks ^b^ |
| --- | --- | --- | --- | --- | --- |
| Study settings |  | 15 | 128 | 2/2 | 0/2 |
| Posterior probability | 60% | 15 | 128 | 2/2 | 0/2 |
|  | *80%* | *15* | *128* | *2/2* | *0/2* |
|  | 95% | 14 | 125 | 2/2 | 0/2 |
| Number of isolates | *3* | *15* | *128* | *2/2* | *0/2* |
|  | 5 | 9 | 107 | 2/2 | 0/2 |
| Proportion local | 0.5 | 28 | 221 | 2/2 | 1/2 |
|  | *0.67* | *15* | *128* | *2/2* | *0/2* |
|  | 0.75 | 15 | 128 | 2/2 | 0/2 |
| Duration | 6 months | 15 | 128 | 2/2 | 0/2 |
|  | *1 year* | *15* | *128* | *2/2* | *0/2* |
| SNP threshold | 50 | 14 | 125 | 2/2 | 0/2 |
|  | 75 | 15 | 128 | 2/2 | 0/2 |
|  | *100* | *15* | *128* | *2/2* | *0/2* |
|  | 150 | 15 | 128 | 2/2 | 0/2 |

^a^ Italicized values are those used in the analysis. When testing alternate values for a parameter, these values are held constant for all other parameters.

^b^ Four resolved outbreaks were selected for testing, two with a known Minnesota source and two with a known non-Minnesota source.

42.	Didelot, X. & Parkhill, J. A scalable analytical approach from bacterial genomes to epidemiology. *Philos Trans R Soc Lond B Biol Sci* **377**, 20210246.
